# Pollutants corrupt resilience pathways of aging in the nematode *C. elegans*

**DOI:** 10.1101/2021.09.28.461818

**Authors:** Andrea Scharf, Annette Limke, Karl-Heinz Guehrs, Anna von Mikecz

**Affiliations:** IUF – Leibniz Research Institute for Environmental Medicine at the Heinrich-Heine-University Duesseldorf, Duesseldorf, Germany; Department of Developmental Biology, Washington University School of Medicine, 660 S Euclid Ave, St. Louis, MO 63110; CF Proteomics, FLI-Leibniz-Institute on Aging, Fritz-Lipman-Institute e.V., Jena, Germany

## Abstract

Delaying aging while prolonging health and lifespan is a major goal in aging research. While many resources have been allocated to find positive interventions with promising results, negative interventions such as pollution and their accelerating effect on age-related degeneration and disease have been mostly neglected. Here, we used the short-lived model organism *C. elegans* to analyze whether two candidate pollutants interfere with positive interventions by corrupting general aging pathways. We took advantage of the immense data sets describing the age-related remodeling of the proteome including increased protein insolubilities to complement our analysis. We show that the emergent pollutant silica nanoparticles (NP) and the classic xenobiotic inorganic mercury reduce lifespan and cause a premature protein aggregation phenotype. Silica NPs rescaled the longevity effect of genetic interventions targeting the IGF-1/insulin-like signaling pathway. Comparative mass spectrometry revealed that increased insolubility of proteins with important functions in proteostasis is a shared phenotype of intrinsic- and pollution-induced aging supporting the hypothesis that proteostasis is a central resilience pathway controlling lifespan and aging. The presented data demonstrate that pollutants corrupt intrinsic aging pathways, which results in premature aging phenotypes. Reducing pollution is therefore an important step to increase healthy aging and prolong life expectancies on a population level in humans and animals.

## Introduction

Aging is a common risk factor for morbidity and mortality in modern human society. In order to prolong life and reduce age-related diseases, millions of dollars are invested into understanding aging and developing effective therapeutics (Partridge et al., 2018). However, these efforts have been made without the consideration of another challenge: global pollution. Since genetic and environmental factors contribute to aging, age-related changes and death (Kenyon, 2010; Sorrentino et al., 2014; Vermeulen et al., 2020), environmental pollution is a likely compounding factor on healthy aging. The Lancet Commission on Pollution and Health reported that pollution is the number one cause for environmentally-induced diseases and premature death (Landrigan et al., 2018). Air, water, and soil pollution caused an estimated 9 million premature deaths worldwide in 2015 (Forouzanfar et al., 2016). The actual number of deaths may be even higher due to many emerging chemicals such as nanoparticles (NPs) or endocrine disruptors that have not yet been studied for their effects. Another neglected aspect in the evaluation of pollution costs, is a pollution-induced decline in quality of life or health span that is not directly linked to a specific disease or death. In summary, pollution is an understudied, negative aging intervention that counteracts healthy aging and reduces life expectancies.

To successfully promote a longer health- and lifespan in humans, a comprehensive analysis of the negative interventions from the pollutome, which includes all pollutants and sources of exposure to them, would be required in addition to the development of therapeutic strategies for anti-aging interventions. However, the large number of pollutants in the pollutome limits this analysis in the human population. In addition, pollutants reflect manufacturing trends over time; therefore, old and emergent pollutants affect the overall composition of the pollutome. For example, inorganic mercury (iHg) represents an ‘old’ pollutant since its concentrations in oceans and soils have been highly elevated due to mining and fossil-fuel combustion for at least 150 years (Streets et al., 2011). Environmental mercury peaked around 1900 due to the North American Gold/Silver Rush and rose again since 1950 with the rise in coal combustion and artisanal gold production (Streets et al., 2011). Since 1950, the chemical industry alone has added more than 150,000 newly synthesized chemicals (Landrigan et al., 2018), and most of these are not systematically analyzed nor are their biological interactions known. Among these, industrial nanoparticles (e.g., silica NPs) experienced mass production at the beginning of the 21st century highlighted by the announcement of the National Nanotechnology Initiative in the United States (Roco, 2003).

The number and compositional complexity of pollutants complicates strategies that seek to understand the effects of the pollutome on human aging. A more feasible strategy is to take advantage of short-lived model organisms to analyze prominent representative ‘old versus emerging’ pollutants on general aging resilience pathways impacted by negative and positive interventions.

*Caenorhabditis elegans* is a free-living nematode roundworm that thrives in microbe-rich habitats on rotting plant material. Its short life cycle has made *C. elegans* to one of the super model organisms of current biomedicine especially aging research (Tissenbaum, 2015). Under abundant growth conditions, a sequence of four larval stages is completed within approximately 2.5 days followed by a hermaphrodite adult stage lasting 2 to 3 weeks (Corsi et al., 2015). A plethora of genes regulate aging of the adult worm, including the insulin/IGF-1-like signaling pathway identified in long-lived *daf-2* (insulin-like growth factor 1, IGF-1) and short-lived *daf-16* (FOXO transcription factor) mutants (Kenyon et al., 1993; Kenyon, 2010). The daf-2 pathway influences the longevity of adult worms as well as aging processes that manifest through the degeneration of neural connectivity, reduced neural signaling, amyloid protein aggregation and acceleration of neuromuscular defects (Collins et al., 2008; Kenyon, 2010; Walther et al., 2015). Notably, the genes of the aging pathways are largely conserved between *C. elegans, Drosophila melanogaster*, mice, and humans (Kaletta and Hengartner, 2006; Kenyon, 2010).

The representative pollutants for ‘old versus emerging’ pollutants iHg and silica NPs are both neurotoxic/neurodegenerative and have the potential to induce amyloid protein aggregation in human cells and *C. elegans* (Arnhold et al, 2015, Scharf et al., 2016). Since it is well established that widespread protein aggregation is a hallmark of aging (David et al., 2010; Hipp et al., 2019), the proteostasis network represents a possible candidate pathway in which to test the effects of pollutants on aging. Proteostasis shapes the proteome through ribosome-associated quality control, chaperone-mediated folding, transporters, and degradation processes via proteasome, lysosomes, and autophagy. Further, proteostasis mechanisms can be modulated by genetic intervention to increase health- and lifespan (Morimoto, 2020).

Here, we demonstrate that the pollutants silica NPs and iHg induce a premature aging phenotype by corrupting resilience pathways that normally preserve protein homeostasis and drive longevity. We show that the tested pollutants not only significantly reduce the lifespan of wild-type *C. elegans*, but also that silica NPs reverse the genetic lifespan resilience effects of *daf-2* mutants. iHg induced proteotoxic stress and characteristics of aggregome network including components of protein folding, proteolysis, and stress response pathways. By comparative mass spectrometry of age- and pollutant-induced aggregomes, we identified a shared aggregation phenotype with an enrichment for proteins involved in protein homeostasis independent of protein abundance. Furthermore, we identified nine super-aggregation proteins with links to human diseases that became insoluble during intrinsic-as well as pollutant-induced aging. These results demonstrate that pollutants negatively affect well described aging prevention pathways and that effective pollution control and reduction will be an important strategy to fight aging and premature death worldwide.

## Results/Discussion

### Silica NPs and inorganic mercury reduce lifespan

To characterize the impact of pollutants on resilience pathways of aging, we analyzed how two widely distributed pollutants, 50 nm silica nanoparticles (NPs) (von Mikecz, 2018) and iHg, affect the lifespan of *C. elegans*. To determine lifespan changes, *C. elegans* hermaphrodites were treated with different concentrations of silica NPs or iHg from the L4/young adult stage in S-Medium with *E. coli* and FUdR at 20°C (Figure 1, Table S1). While low concentrations of silica NPs did not show any significant changes in lifespan, 1.25 mg/mL silica NPs decreased the mean and maximum lifespan by 35% and 24% respectively (Figure 1A). In contrast, iHg decreased lifespans in a concentration-dependent manner with 10, 25, and 50 μM iHg reducing lifespans by 14%, 63%, and 86%, respectively (Figure 1B, Table S1). Maximum lifespans after iHg exposure were decreased by 16%, 62%, and 78%, respectively (Figure 1B, Table S1).

**Figure 1.**
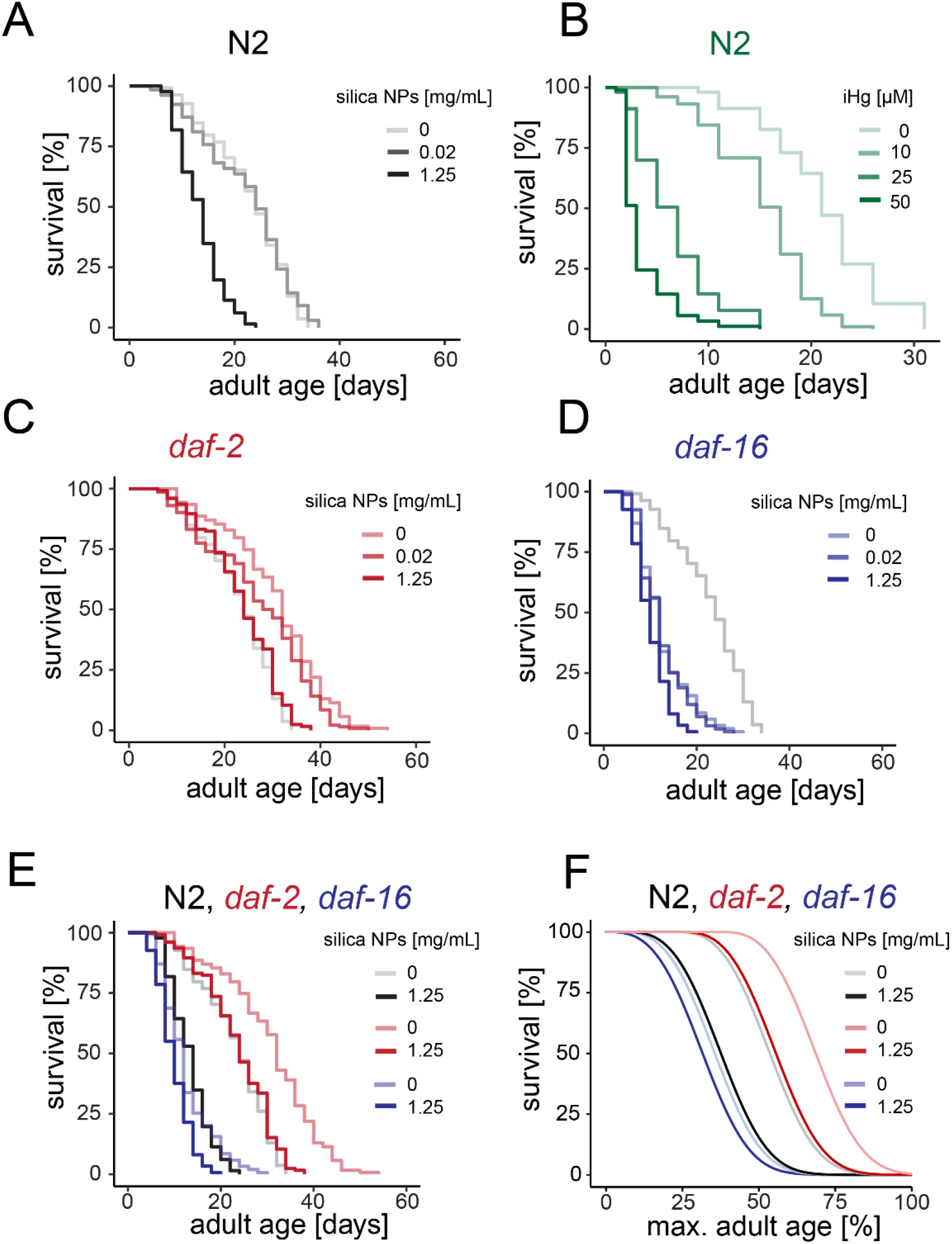
Environmental stressors rescale *C. elegans* lifespans. (A-F) Animals were cultured in S-medium with OP50 as food source and supplemented with FUDR to prevent larval hatching. (A) Representative survival curves for wild type (N2) exposed to silica NPs from adult day 1 as indicated. See table S1 for summary statistics. (B) Representative survival curves for wild type N2 exposed to iHg from adult day 1 as indicated. See table S1 for summary statistics. (C) Representative survival curves for *daf-2* (e1368) exposed to silica NPs from adult day 1 as indicated. See table S1 for summary statistics. (D) Representative survival curves for *daf-16* (mu86) exposed to silica NPs from adult day 1 as indicated. See table S1 for summary statistics. (E) Representative survival curves for wild type N2, *daf-2* (e1368), *daf-16* (mu86) exposed to silica NPs from adult day 1 as indicated. See table S1 for summary statistics. (F) The blue, grayscale and red (Weibull) lines represent maximum-likelihood estimation (MLE) fits of the parametric model using data up to median survival.

### Silica NPs reverse genetic lifespan resilience effects

Next, we investigated whether aging interventions protect against the lifespan shortening effects of silica NPs using *C. elegans* strains carrying mutations in the IGF signaling pathway. Mutations in the *daf-2*/insulin/IGF-1-receptor gene extend lifespan via activation of the DAF-16/FOXO transcription factor-mediated protection gene program, whereas *daf-16(lf)* strains show reduced lifespans compared to wild-type (Kenyon et al., 1993; Piechulek & von Mikecz, 2018). We chose this genetic intervention due to its effective lifespan extension phenotype and potential to prevent protein aggregation (David et al., 2010), since silica NPs induce premature protein aggregation in *C. elegans* (Scharf et al., 2013). Silica NP treatment significantly decreased the mean and maximum lifespan of *daf-2(e1368)* mutant animals by 23% and 30% (Figure 1C, Table S1). Similarly, *daf-16(mu86)* loss-of-function mutant animals (Lin et al., 1997) exhibited a 21% shorter mean and a 33% shorter maximum lifespan when treated with silica NPs (Figure 1D, Table S1). Although silica NPs decreased the lifespans of wild-type and both mutant strains, we observed a temporal rescaling dependent on DAF-16 activity (Figure 1E, F, Table S1). Silica NP exposure shifted the lifespan distribution of *daf-2(lf) mutants* back to wild-type and of wild-type animals back to *daf-16(lf)* mutants. This result indicates that DAF-16 activity protects *C. elegans* from silica NP-induced premature death and restores wild-type lifespan. However, these findings also indicate that silica NPs interfere with the longevity effect and reduce the lifespan of daf-2 mutants to wild-type. In conclusion, silica NPs diminished any effect of the positive genetic intervention on prolonging the lifespan.

### iHg induces proteotoxic stress

Temporal scaling of lifespan distributions is a feature of most aging interventions that is based on a shared organismal state also called resilience or vitality (Stroustrup et al., 2016). Therefore, if we consider pollutants as negative aging interventions, they would also affect aging by influencing the same organismal state or resilience pathway. Proteostasis is a possible candidate for such a resilience pathway given that (1) disruption of proteostasis is a hallmark of intrinsic aging that manifests as the accumulation of protein aggregates (Hipp et al., 2019), and (2) several studies have shown that pollutants and xenobiotics can induce protein fibrillation and disrupt proteostasis (Uversky et al., 2001; Chen et al., 2008; Arnhold et al., 2015; Scharf et al., 2016). Furthermore, xenobiotic exposure can be linked to neurodegenerative diseases such as Parkinson’s or Alzheimer’s disease and can induce pathogenic protein aggregation of amyloid-like assemblies containing several hundred proteins as well as nonproteinaceous components including metals (Uversky et al., 2001; Gerhardsson et al., 2008; Meleleo et al., 2020; Lashuel, 2021).

We predicted that if proteostasis is the underlying resilience pathway, silica NPs and iHg-exposed *C. elegans* would share features of the protein aggregation phenotype with aged *C. elegans*. To investigate this, we first used a transgenic *C. elegans* model for proteotoxic stress in muscle cells that allows the visualization of age-related induction of polyglutamine (polyQ) fibrillation (Morley et al., 2002). This strain expresses homopolymeric repeats of 35 glutamines fused to yellow fluorescent protein (Q35::YFP) in muscle cells of the body wall. Young Q35::YFP *C. elegans* showed a smooth fluorescent pattern along the muscle fibers which changes into an irregularly dotted pattern of polyQ aggregates with age (Morley et al., 2002). We exposed Q35 animals on adult day 1 to 50 μM iHg or left them untreated (H_2_O) for 24 hours. Untreated Q35 *C. elegans* showed a mostly smooth distribution of yellow fluorescence along the body wall muscles on adult day 2 (Figure 2A,B, S1), consistent with previously published observation (Morley et al., 2002). In contrast, exposure to iHg prematurely induced the dotted pattern of aggregated polyQ typical for animals older than adult day 3 (Figure 2A,B, S1). Quantification of the fluorescent polyQ aggregates showed a significant 1.9-fold increase in the number of aggregates after iHg exposure. This result indicates that iHg induces proteotoxic stress prematurely in young *C. elegans*, which is supported by the previous observations that iHg induces fibrillation of endogenous proteins in the nucleolus of intestinal cells in *C. elegans* as well as causes aggregation of nuclear proteins in human cells (Table S2). Notably, silica NPs induced the same age-related aggregation phenotype as we showed in a previous study (Table S2) (Scharf et al., 2016). In summary, the two candidate pollutants iHg and silica NPs induce widespread protein aggregation in young *C. elegans* that manifests on the local and global level representing a premature aggregation phenotype that normally develops during the intrinsic aging process.

**Figure 2.**
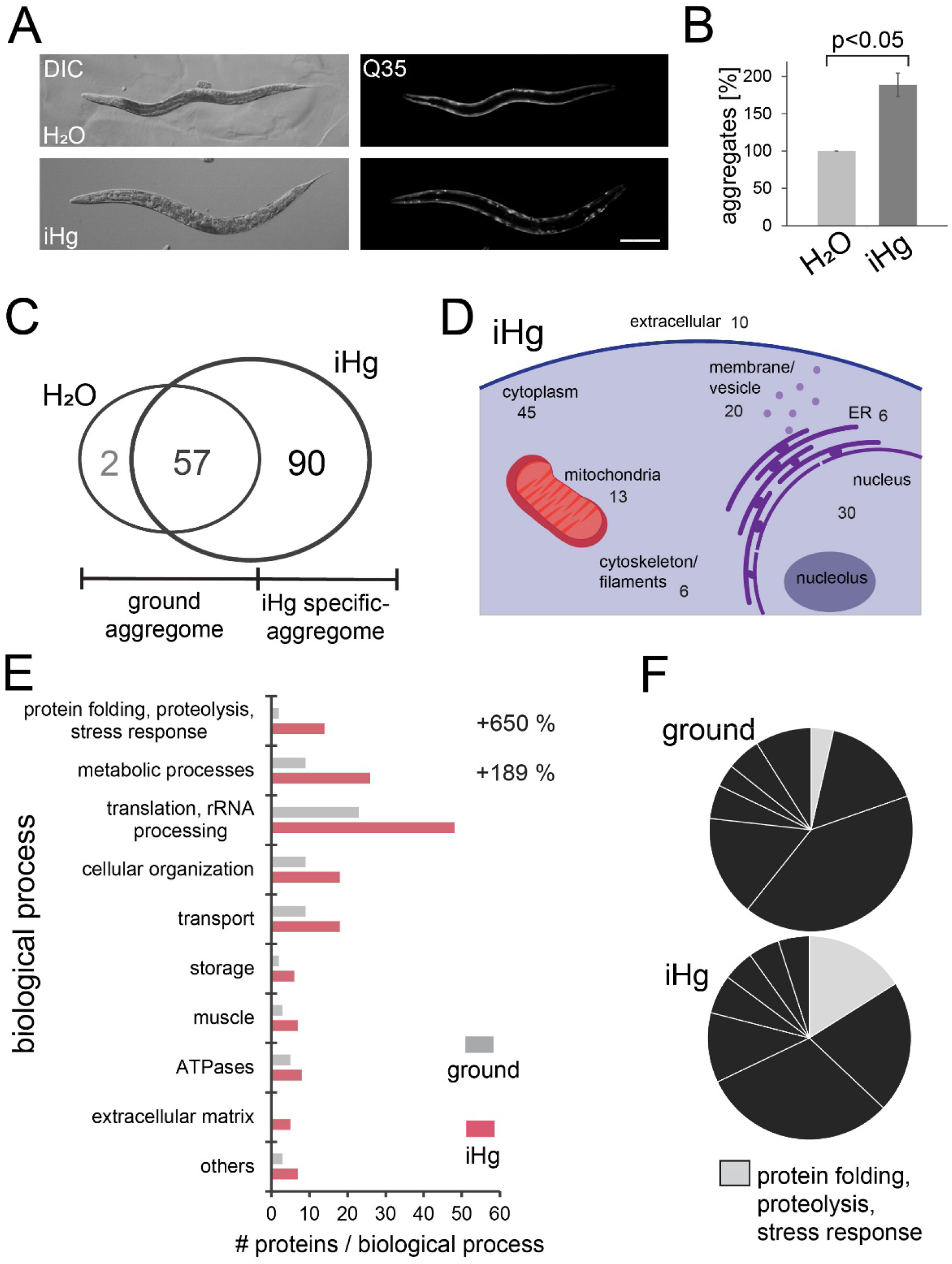
Inorganic mercury accelerates protein fibrillation in adult *C. elegans*. (A) Representative fluorescent micrographs of 2-day old, adult hermaphrodites of a *C. elegans* proteotoxic stress model that expresses homopolymeric repeats of 35 glutamines fused to yellow fluorescent protein (Q35::YFP) in body wall muscle cells (Morley et al., 2002). Hermaphrodites were mock-treated or exposed to 50 μM iHg for 24 hours. Bar, 100 μM. (B) Respective quantification of the increase in Q35-YFP aggregates per hermaphrodite compared to control. Values represent +/- SD of three independent experiments with n>90 for each treatment. Student’s *t* test *p<0*.*05* for H_2_O versus iHg. (C) Venn-diagram of the insoluble protein fraction of 24-hour mock-treated or iHg-exposed wild type *C. elegans*. We defined proteins that aggregate specifically in iHg-exposed *C. elegans* as iHg-specific aggregome and all other SDS-insoluble proteins as ground state aggregome respectively. (D) Cellular localization of iHg-induced insoluble proteins. The numbers indicate the number of identified candidate proteins associated with each cellular subcellular compartment according data mining of the PANTHER database (Mi et al., 2013), UniProt (Bateman et al., 2021), and WormBase (Harris et al., 2020). (E) GO analysis of the ground and iHg-induced aggregome. Each identified insoluble protein was categorized according to its biological process classified using the PANTHER database, UniProt, WormBase, and literature. The graph presents the absolute number of identified SDS-insoluble proteins per biological process category. The percentage indicates the change in the number of identified proteins in the respective functional group in the ground versus the iHg-induced aggregome network. (F) The functional group protein folding, proteolysis, and stress responds shows the highest increase in insoluble proteins in iHg-exposed *C. elegans* as illustrated in the pie chart.

### iHg-induced aggregome network

In a previous study, we identified the components of the silica NP-induced aggregome network in *C. elegans* via filter retardation assay and mass spectrometry (Scharf et al., 2016). To now characterize the iHg-induced aggregation network in order to compare the components to the silica NPs-induced aggregome network, wild-type day 1 adults were either left untreated (H_2_O) or treated with 50 μM iHg for 24 hours, lysed, and the SDS-insoluble protein fraction isolated via filter retardation assay (Scheme, Figure S2, (Scherzinger et al., 1997). This method specifically traps SDS-insoluble high molecular weight protein aggregates, also referred to as amyloid-like protein aggregates. The soluble fraction, including less stable amorphous protein aggregates, passes through the 0.2 μm pores of cellulose acetate membranes, whereas the amyloid-like proteins are trapped (Wanker et al., 1999). In order to identify the amyloid-like proteins, we eluted and analyzed the trapped SDS-insoluble protein fraction via mass spectrometry, gene ontology (GO) analysis, and data mining. We identified 90 proteins that aggregated specifically in iHg-exposed *C. elegans*, whereas 57 proteins aggregated independently of iHg exposure (Figure 2C, Table S3). We defined these two groups of SDS-insoluble proteins as mercury-specific aggregome and as ground state aggregome, respectively. Although proteins of the iHg-specific aggregome localized throughout the cell, proteins were mostly associated with the cytoplasm, nucleus, and membranes (Figure 2D). Interestingly, the nucleus showed the largest fraction of different insoluble proteins in the ground state and after iHg exposure. The insoluble fraction included many ribosomal proteins, but also prominent nucleolar proteins such as fibrillarin (FIB-1) (Table S3). GO analysis revealed that the iHg-induced aggregome network is enriched with proteins related to the biological functions of “metabolic processes” (17 out of 90 proteins) and “translation and RNA processing” (25 out of 90 proteins) (Figure 2E). Furthermore, proteins with the biological function of “proteolysis and stress-response” showed the largest increase with 650% after iHg exposure (Figure 2E,F). The two functional groups “translation and RNA processing” and “proteolysis and stress-response” can be combined as functional group “proteostasis”, accounting for over 42% of proteins identified in the iHg-specific aggregome.

Comparison with the previously described silica NP-induced aggregome network (Scharf et al., 2016) revealed 20 shared proteins that fibrillate after exposure to iHg and silica NPs (Figure 5, Table S3). In addition, both aggregome networks showed the largest enrichment of proteins with the biological functions “metabolic processes”, “translation and RNA processing”, and “proteolysis and stress-response”. Therefore, proteins that drive protein homeostasis (“translation and RNA processing” and “proteolysis and stress response” combined) showed the largest increase after exposure to silica NPs and iHg. Specifically, the chaperones HSP-6 and HSP-60 as well as the proteasome subunits RPN-2 and RPN-3 were shared components of both aggregomes. These proteins are essential for maintaining the proteostasis and likely sequester to microenvironments in the cell to dissolve protein aggregations. In addition, many proteins that are essential for the first steps of proteostasis such as protein translation were also part of the aggregomes. While chaperones and proteasomes may be functional components of the amyloid-like protein aggregates (Chen et al., 2008), proteins necessary for protein synthesis are most likely dysfunctional in their amyloid-like protein state. As a consequence, fine-tuned cellular and organismal proteostasis is disturbed. This result supports the hypothesis of proteostasis as a central resilience pathway controlling lifespan and aging.

### Age-induced aggregome-network

Next, we asked whether iHg and silica NPs (1) accelerate age-related aggregation of proteins or (2) induce the insolubility of a specific set of proteins that are different from the age-induced aggregome. In order to answer this question, we analyzed age-induced insoluble protein data sets from the literature. The Kenyon lab identified 461 proteins with an age-dependent shift to an insoluble state using two temperature sensitive sterile strains: the gonad-less mutant *gon-2* and the germline-deficient mutant *glp-1* at 25°C (David et al., 2010). The Lithgow and Hughes labs identified 203 proteins that shift age-dependently to an insoluble state by using the temperature sensitive sterile strain TJ1060 (*spe-9(hc88)I; fer-15(b26)II*) (Reis-Rodrigues et al., 2012). These two datasets shared 117 proteins that became insoluble in aged *C. elegans* (Figure 3E, Table S4,5). Both approaches used centrifugation-based separation of the insoluble and soluble protein fractions with SDS-containing buffers and resuspension with 70% formic acid. We reasoned that by using the amyloid-specific filter trap assay (Scherzinger et al., 1997) to analyze the aged aggregome network in wild-type *C. elegans*, we would add a dataset containing proteins that form age-dependent amyloid-like aggregates. Therefore, wild-type (N2) *C. elegans* were cultured on agar plates supplemented with FUdR to prevent egg hatching and lysed at adult day 1 (young) and adult day 12 (old). The SDS-insoluble protein fractions were isolated via filter retardation assay (Figure S2, Scherzinger et al., 1997) and analyzed by mass spectrometry. We identified 79 proteins that aggregate specifically with old age, whereas 31 proteins are already insoluble in young animals (Figure 3A, Table S6). We defined these two groups as old age-specific aggregome and as ground aggregome respectively. Over 65% of the identified proteins of the old age-specific aggregome were enriched in the insoluble protein fractions reported in the David et al., 2010 and Reis-Rodriguez et al., 2012 data sets. All three data sets shared 22 old age-induced insoluble proteins, seven of which play a role in proteostasis (Figure 3E, Table S4,5). Thus, comparative mass spectrometry based on different biochemical preparations clearly provides an added value by further clarification of the cellular pathways. Here, the comparative dissection of age-induced protein aggregates reveals proteostasis as a specific pathway of amyloid formation.

**Figure 3.**
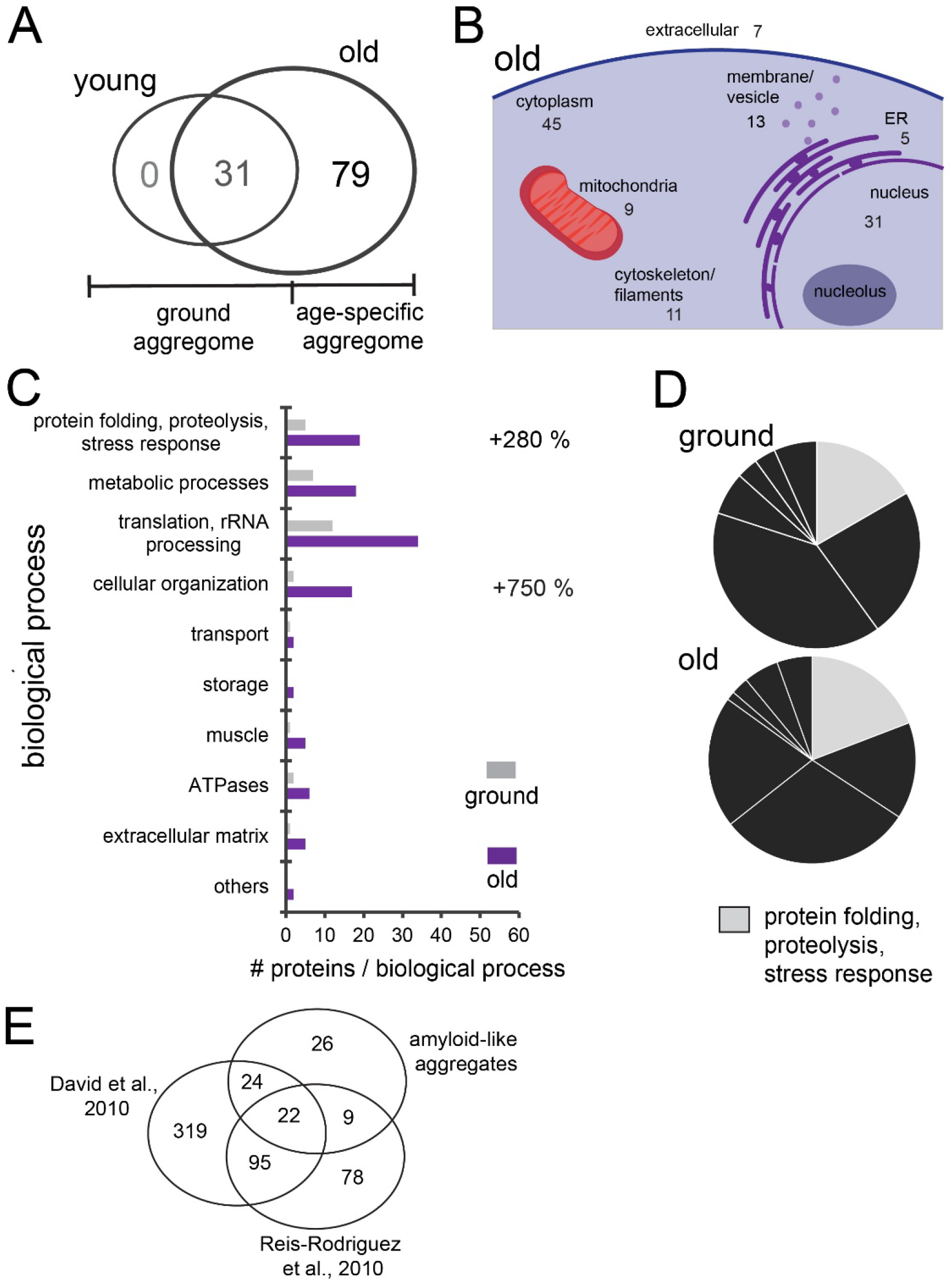
Age-related amyloid protein fibrillation in *C. elegans*. (A) Venn-diagram of the insoluble protein fraction of 1- (young) and 12-day (old) adult wild type *C. elegans*. We defined proteins that aggregate specifically in 12-day old adult *C. elegans* as age-specific aggregome and all other SDS-insoluble proteins as ground state aggregome respectively. (B) Cellular localization of age-induced insoluble proteins. The numbers indicate the number of identified candidate proteins associated with each cellular subcellular compartment according data mining of the PANTHER database (Mi et al., 2013), UniProt (Bateman et al., 2021), and WormBase (Harris et al., 2020). (C) GO analysis of the ground and age-induced aggregome. Each identified insoluble protein was categorized according to its biological process classified using the PANTHER database, UniProt, WormBase, and literature. The graph presents the absolute number of identified SDS-insoluble proteins per biological process category. The percentage indicates the change in the number of identified proteins in the respective functional group in the ground versus the age-induced aggregome network. (D) The change of the functional group protein folding, proteolysis, and stress responds in young versus old *C. elegans* is illustrated in the pie chart. (E) Comparative mass spectrometry based on different biochemical preparations: Venn diagram of insoluble proteins identified in David et al., 2010 and Reis-Rodrigues et al., 2012 compared to the amyloid-like protein aggregates identified in this study.

### Pollutant- and old age-induced aggregation phenotype

Next, we compared the old age-induced aggregome with the iHg-induced aggregome network and observed that proteins of both aggregomes localize throughout the cell. In agreement with the iHg-specific aggregome, the cytoplasm, nucleus, and membranes of the age-induced aggregome network showed the highest amount of age-induced insoluble proteins (Figure 3B). GO-analysis revealed that the old age-induced aggregome network was also enriched for proteins related to the biological functions of “proteolysis and stress-response”, “metabolic processes”, and “translation and RNA processing” (Figure 3C). In contrast to the iHg- and silica NP-induced aggregome networks, proteins with the biological functions of “cellular organization” showed the largest increase with 750% of proteins in the SDS-insoluble amyloid-like protein fraction in old *C. elegans* compared to the ground state. While structural proteins such as the intermediate filament IFB-2 (human ortholog lamin B2) or IFC-2 (human orthologs include keratin 72,74,83) are not affected by xenobiotic-induced premature aging, they are an intrinsic part of the old age-induced aggregome network.

One possible explanation is that these structural proteins aggregate as a consequence of the age-related reduced capacity of chaperones and clearance machinery. The decline of the proteostasis network due to reduced expression (Ben-Zvi et al., 2009) and increased sequestration of its components to protein aggregates with age causes an increasing solubility imbalance in the organism highlighted by the insolubility of structural proteins in the aged-induced aggregome. In agreement, IFB-2 was identified as one of 10 age-dysregulated proteasomal targets, which mislocalizes into cytosolic protein aggregates linked to loss of intestinal integrity (Koyuncu et al., 2021). Further, knockdown of *ifb-2* in adult *C. elegans* results in an increased lifespan and better intestinal health (Koyuncu et al., 2021). Future studies that include longitudinal experiments could time stamp the appearance of different functional protein groups in the insoluble fraction and allow construction of an insolubility time line.

We also compared the structural features of the proteins in old age-, iHg-, and silica NPs-induced aggregome networks (Figure 4). All three aggregome networks were enriched with proteins containing beta strands, cross-links, helices, nucleotide binding capacities and repeats. Only a few features were specific for a single aggregome. The old age-induced aggregome showed more DNA-binding proteins, whereas the xenobiotic-induced aggregomes exhibited an overrepresentation of zinc finger-containing proteins. Proteins with metal binding domains were only enriched in the iHg-induced aggregome.

**Figure 4.**
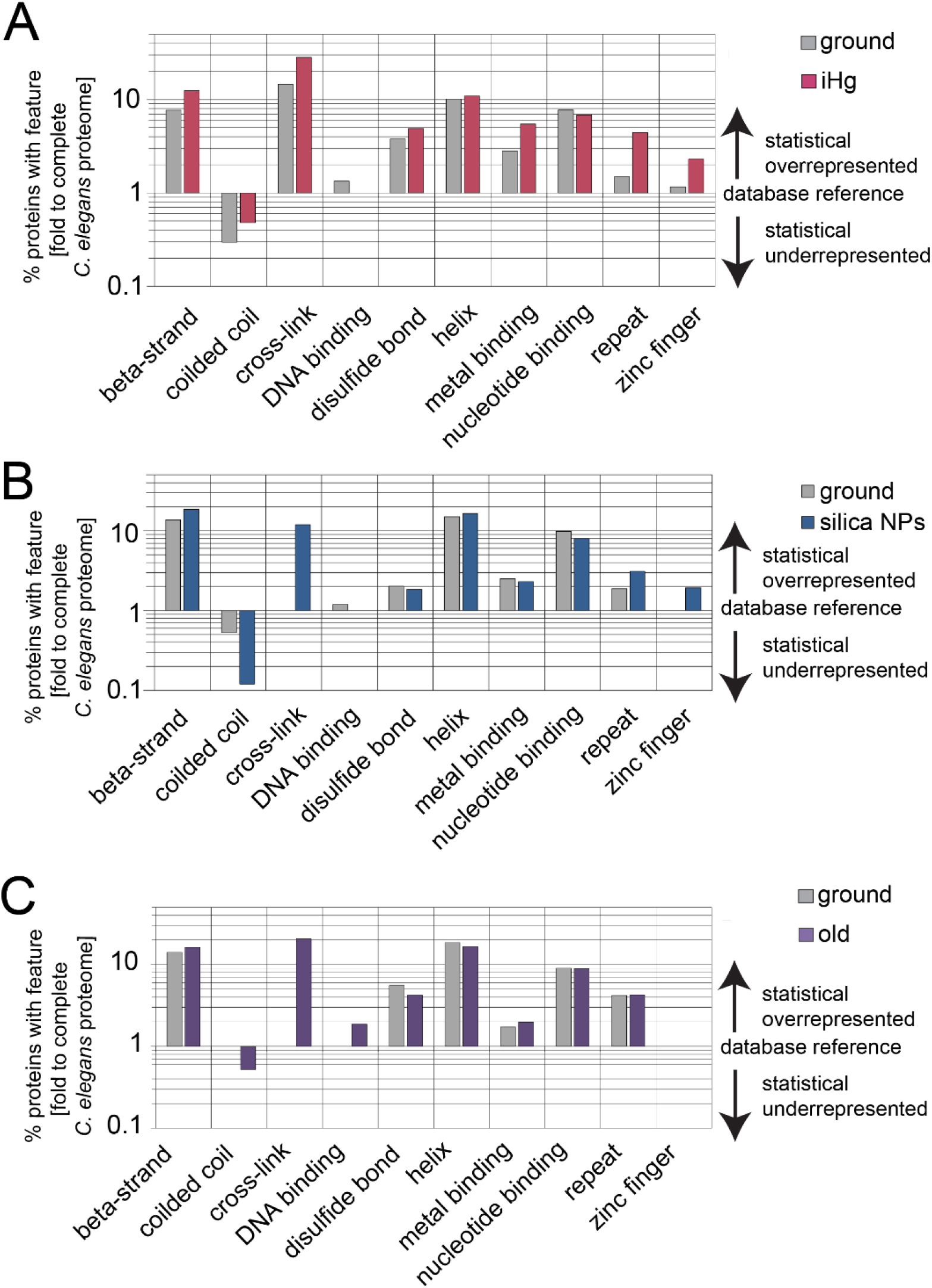
Shared structural protein feature of aggregome networks. Analysis of structural protein features of the (A) iHg-, (B) silica NPs- (Scharf et al., 2016), and (C) age-induced aggregomes based on the UniProt database (Bateman et al., 2021). The identified fibrillated proteins show a statistical overrepresentation in beta strands, cross links, helices, and nucleotide binding. Metal binding is only overrepresented in the iHG-induced aggregome, and zinc fingers are absent from the age-induced aggregome.

Overall, all three aggregome networks showed many similarities including protein features and functional groups. Proteins that drive the protein homeostasis were the most enriched proteins in the old age-, iHg-, and silica NP-induced aggregome networks (Table S3,6, Figure 2,3,S3, S4 (Scharf et al., 2016)). This result supports the hypothesis that proteostasis represents a central resilience pathway, controlling lifespan. We showed that *C. elegans* exposed to iHg- and silica NPs prematurely induce an old age aggregome network, consistent with the idea that pollutants accelerate aging by corrupting the respective resilience pathways of intrinsic aging.

### Identification of super aggregation proteins

Given that the two pollutants studied, iHg and silica NPs, prematurely induce the same protein aggregation phenotype as intrinsic aging, we would expect that the same proteins segregate into the aggregome network. Consistent with this, we found nine super aggregation proteins that were present in the iHg-, silica NP-, and age-induced aggregome networks (Figure 5, Table 1), suggesting the three aggregome networks are to more than 10% identical. The silica NPs- and iHg-aggregome networks shared 11 proteins, whereas 14 were shared between the iHg-induced and old aggregome network and 12 proteins were shared between the silica NPs-induced and the old aggregome network.

**Figure 5.**
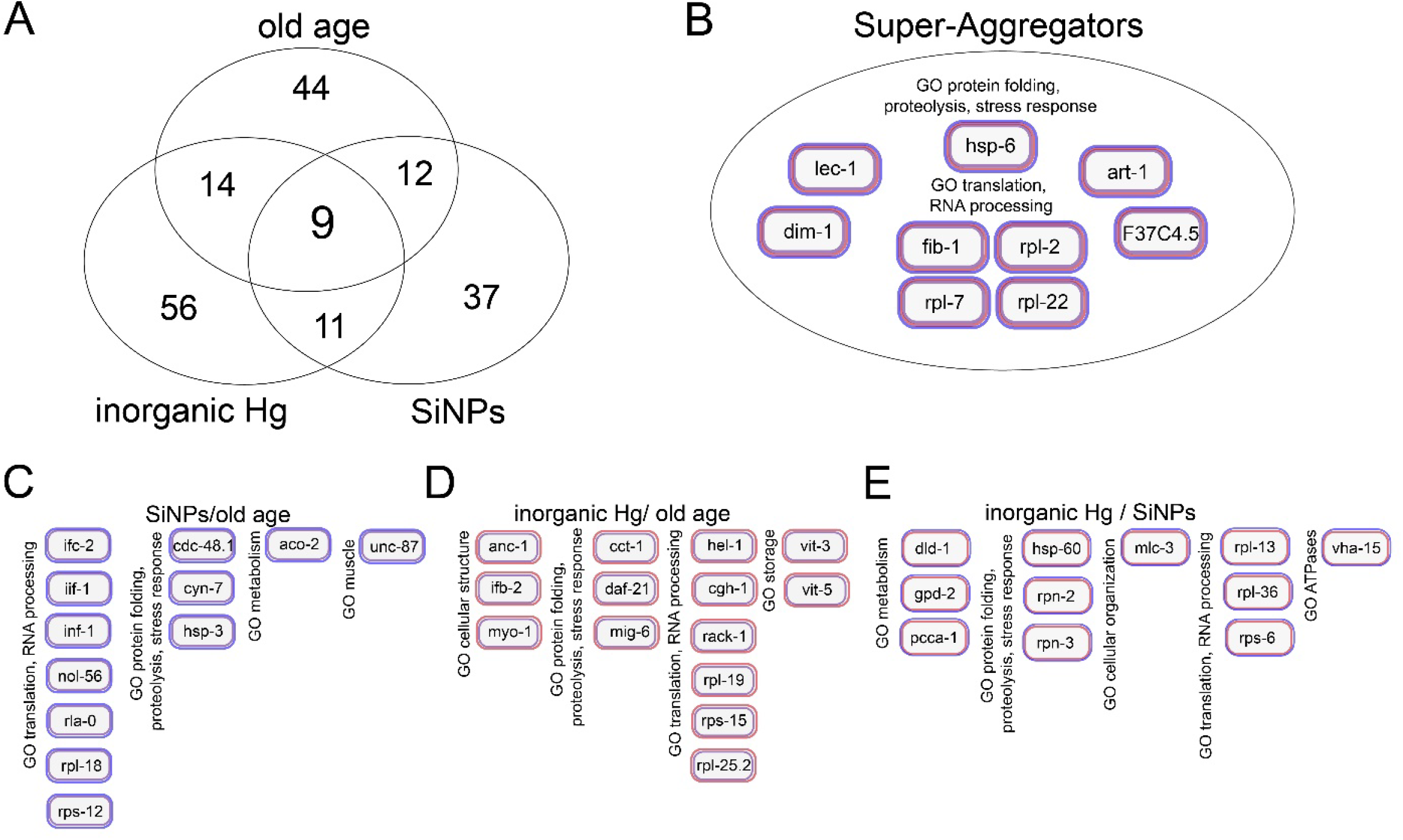
Identification of 9 super aggregators shared between aggregomes. (A) Venn diagram of insoluble proteins of the iHg-, silica NPs- (Scharf et al., 2016), and the age-induced aggregome. (B) All three aggregomes share 9 super aggregator proteins: HSP-6, FIB-1, RPL-2, RPL-7, RPL-22, ART-1, LEC-1, DIM-1, and F37C4.5. Shared identified insoluble proteins between (C) the silica NPs- and the age-induced aggregome, (D) iHg- and the age-induced aggregome, and (E) between the iHg- and silica NPs-induced aggregome.

**Table 1.**
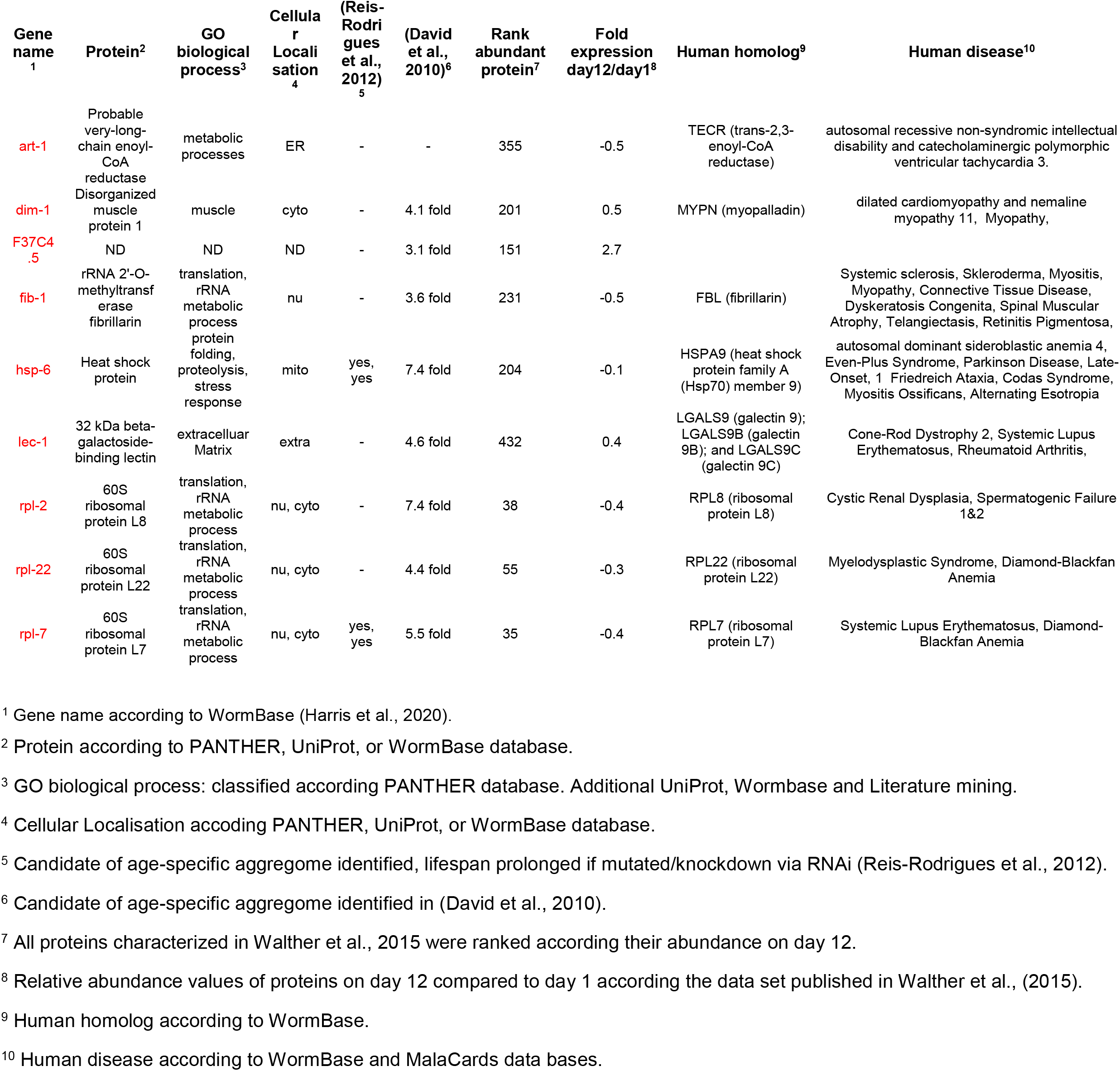
Super Aggregators are identified in the iHg-, silica NPs-, and age-specific aggregome.

An alternative explanation for the identification of proteins with similar functions and even shared proteins in all three aggregomes could be their abundance in the tissues of *C. elegans*. To exclude that the identified proteins are just the highest expressed proteins under the different conditions, we first mined the mass spectrometry dataset from Walther et al., 2015 for changes in abundance levels of proteins in old compared to young N2 *C. elegans* (Table 1). AHCY-1 was the most abundant protein in day-1 *C. elegans* in the data set (Walther et al., 2015) and was identified as insoluble protein in the ground aggregome here. In contrast, RPL-30 was the second most abundant protein in adult day 1 *C. elegans* and 28^th^ most abundant protein in adult day 12 worms (Walther et al., 2015), but was not identified in any aggregome. All nine super aggregation proteins ranked in the first 600 out of nearly 6000 proteins with abundance data in adult day 1 and adult day 12 *C. elegans* (Table 1). Nevertheless, 7 of the 9 proteins are less abundant in 12-day old compared to 1-day old *C. elegans* (Table 1). This trend was confirmed with qPCR for the protein FIB-1 and HSP-60. Both proteins did not show any significant increase in mRNA expression after iHg-treatment, whereas HSP-6 exhibited a 11-fold induction in mRNA expression (Figure S5). Although we concluded that the identified aggregomes do not simply reflect just the most abundant proteins, it is still possible that highly abundant insoluble proteins are more likely detected than less abundant insoluble proteins.

The nine super aggregation proteins were ART-1, the probable very-long-chain enoyl-CoA reductase; HSP-6, a mitochondrial chaperone; FIB-1, the rRNA 2’-O-methyltransferase fibrillarin; DIM-1, the disorganized muscle-protein 1; LEC-1, a 32 kDa beta-galactoside-binding lectin; the unnamed protein F37C4.5; and the three ribosomal proteins RPL7/L7, RPL2/L8, and RPL22/L22 (Table 1). All of these proteins, with the exception of ART-1, were also identified in the age-induced insoluble protein fraction of two temperature sensitive sterile strains: the gonad-less mutant *gon-2* and the germline-deficient mutant *glp-1* at 25°C (David et al., 2010), which validates the significance of these super aggregation proteins in the development of aggregome networks. Furthermore, all proteins, with the exception of F37C4.5 (orthologs are not known), have homologs in humans that are linked to human diseases (Table 1). The majority of these super aggregators are protein homeostasis proteins, which supports the hypothesis of proteostasis as a central resilience pathway controlling aging and lifespan.

Interestingly, experiments with HSP-6 and RPL-7, two proteostasis proteins, showed opposite effects on *C. elegans* lifespan in previous studies: *hsp-6* RNAi reduced and HSP-6 overexpression increased lifespan significantly (Kimura et al., 2007), whereas *rpl-7* RNAi significantly increased the lifespan by more than 16% (Reis-Rodrigues et al., 2012). These results seem to be contradicting, but the functions of these two proteins can explain the different effects on lifespan. RPL-7 is a ribosomal protein that localizes in the nucleolus during ribosomal RNP subunit assembly and in the cytoplasm as part of ribosomes. While RPL-7 is normally abundant in the cell, it shows a 0.4 fold decreased relative abundance in the proteome of old worms compared to young worms (Table 1, Walther et al., 2015). Consequently, the appearance of RPL-7 as a super aggregation protein in induced aggregome networks is not just caused by abundance, although it is still the 35^th^ most abundant protein in old worms (Walther et al., 2015). Ribosomal proteins are aggregation-prone due to unfolded extensions and basic regions that are essential for their interactions with rRNAs (Pillet et al., 2017). It seems likely that such a protein will fibrillate and drive protein aggregation once the proteostasis is misbalanced due to aging or pollutant stressors. Knockdown of *rpl-7* via RNAi in adult animals reduces the abundance of RPL-7 without affecting the animals’ development and thereby decreases also the amount of RPL-7 that has the potential to fibrillate. As a consequence, *rpl-7* RNAi reduces the burden of insoluble proteins and crowding effects in the nucleus/nucleolus (Vecchi et al., 2020), which results in an increased lifespan. In contrast to RPL-7, HSP-6 is a chaperone and part of the protein folding system of the organism (Ruan et al., 2020; Shin et al., 2021). HSP-6 is the mitochondrial HSP-70, highly conserved, and has the ability to prevent aggregation (Bender et al., 2011). The protein is prone to self-aggregation with the important feature that aggregated mHSP70 preserves its chaperone function (Kiraly et al., 2020). Nevertheless, it seems likely that HSP-6 and other chaperones become part of the aggregome networks by binding to increasing amounts of misfolded and fibrillated proteins. Due to their sequestration into protein aggregation, these chaperones are unavailable for their physiological clients to assist them during folding and sustain a healthy proteome, a process called chaperone competition (Yu et al., 2019), creating a viscious cycle. Chaperones are occupied with the increasing pool of insoluble proteins and therefore unavailable for freshly synthesized proteins in need of folding support (Santra et al., 2019). These proteins join the pool of insoluble proteins and make chaperones even more necessary. Escape is impossible due to the dramatic decline of chaperone expression in early adulthood (Ben-Zvi et al., 2009). Consequently, the organism experiences widespread protein aggregation, protein dysfunction, and failure of organs and tissues with age (Santra et al., 2019), which can be accelerated through pollutant exposure.

## Conclusion

The concept of environmentally-induced disease and premature death (Landrigan et al., 2018) is supported in the animal model *C. elegans*. Here, we show that the well-known xenobiotic iHg and the emerging pollutant silica NPs corrupt the same resilience pathways that normally preserve cellular homeostasis and thereby accelerate aging processes (Figure 6). Silica NPs accelerates global protein aggregation (Scharf, Piechulek, & von Mikecz, 2013) and reverse the genetic activation of resilience pathways through loss of function mutations in the *daf-2* gene. The capacity of the proteostasis network to resolve misfolded and damaged proteins declines in early adulthood in *C. elegans* (Ben-Zvi et al., 2009) and causes an age-related increase in protein aggregation manifested in an old age aggregation phenotype. The decline of the proteostasis network is most likely temporally fine-tuned to secure reproduction which peaks on day 2/3 of adulthood under normal condition (Scharf et al., 2021). This conclusion is supported by the observation, that most somatic age-related changes occur after the decline of reproduction (Pickett et al., 2013). Exposure to pollutants such as iHg and silica NPs adds an additional proteotoxic stress to the declining proteostasis network. As a result, the fine-tuned resilience pathway reaches the tipping-point earlier resulting in premature aggregation phenotypes, accelerated somatic decline (Scharf et al., 2013; Piechulek et al., 2019), and premature death of the animal.

**Figure 6.**
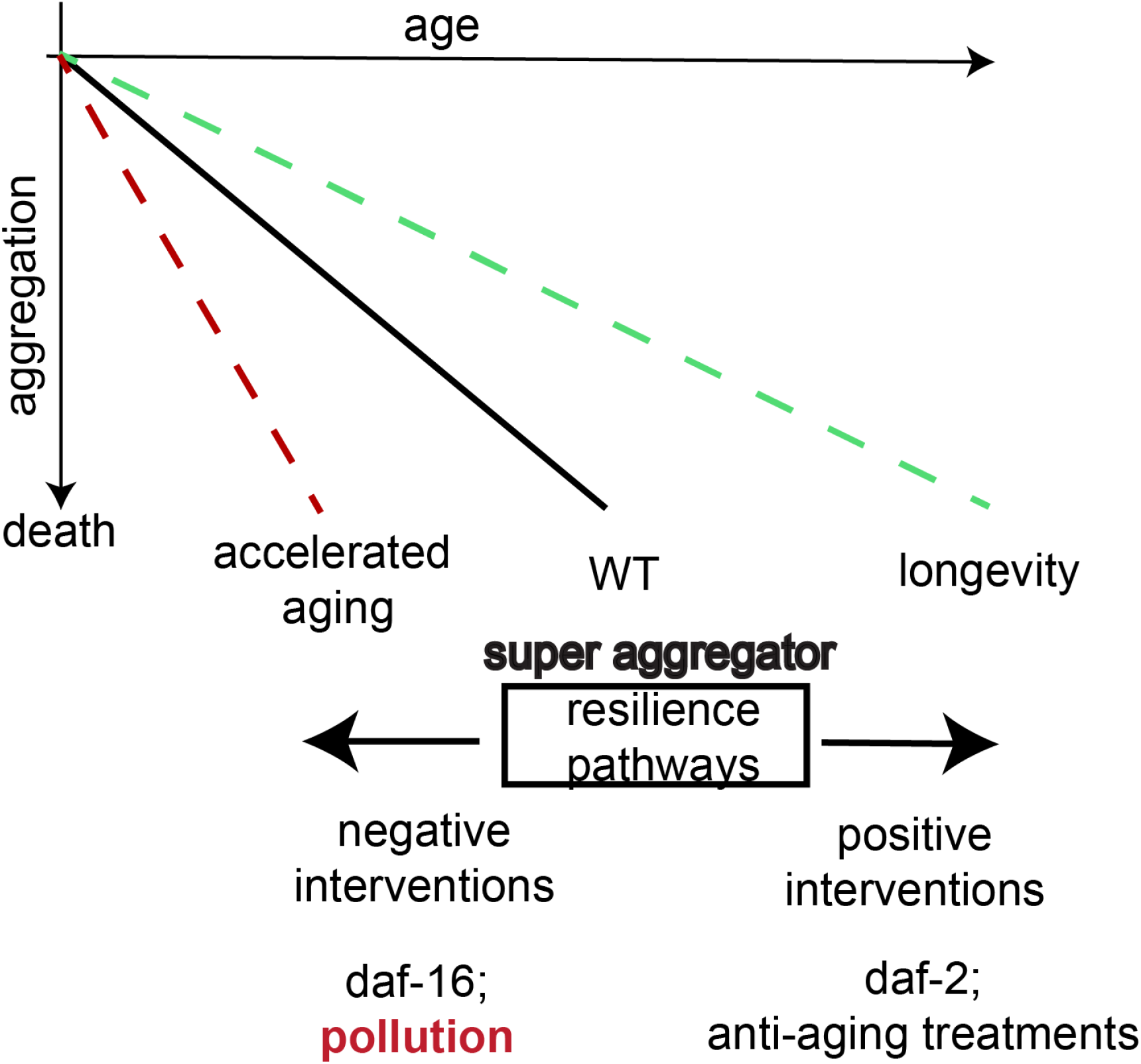
Pollutants hijack resilience pathways of intrinsic aging. The schematic shows how negative interventions and positive interventions change the fibrillation status of an organism and affect its aging rate. We propose that genetic interventions such as *daf-16* and *daf-2* loss of function mutations, anti-aging treatments such as HSP-90 inhibitors, or pollution such as the well-known pollutant iHg as well as the emerging pollutant silica NP affect the same resilience pathways that normally preserve cellular homeostasis and thereby accelerate or delay aging processes. As a conclusion, reducing pollution and controlling the synthesis of new compounds has to be an important part of an effective strategy to support healthy aging.

The identified fibrillated proteins can be categorized in four different groups: (1) functional amyloid, (2) highly unstructured and “sticky”/aggregation-prone proteins, (3) components of the proteostasis network, (4) “victims” of failed proteome maintenance. (1) Functional amyloid proteins can form fibrillar aggregates as part of their executive function. For example, many RNA/DNA interacting proteins contain polyQ repeats that support the dynamic interaction at promoters during gene expression (von Mikecz, 2014). These proteins are sensitive to the crowding status of the cellular environments, and overcrowding can easily shift the balance to dysfunctional fibrillation. Proteins such as FIB-1 and many ribosomal proteins fit into this category. (2) By contrast, many of the identified proteins exhibit unstructured regions or protein features that are aggregation-prone and result in pathological fibrillation in the absence of a functional proteostasis network. Interestingly, HSP-6 and proteasome subunits belong in this category. (3) Components of the proteostasis network such as chaperones or the proteasome bind to fibrillated proteins in order to restore functional folding, therefore, sequestration to local protein aggregation environments is part of their function. (4) Finally “victims” of failed proteome maintenance are proteins that are normally refolded and/or cleared, but remain in their cellular environment due to missing proteostasis network capacities. Proteins such as IFB-1 fall into this category and are more specifically associated with the age induced-aggregome. Proteins of all four groups constitute the amyloid protein fibrillation landscape spanning from facilitation of functional protein interactions to pathological aggregates (von Mikecz, 2009; Lashuel, 2021). Consistent with this, deciphering the proteomes of intrinsic vs pollutant-induced aging has the potential to advance translational research in neurodegenerative diseases where age is the prominent risk factor.

We show that pollutants corrupt resilience pathways of intrinsic aging. As a result, pollutants negatively affect the same prevention pathways of aging that are targeted by possible anti-aging therapeutics such as HSP90 inhibitors (Janssens et al., 2019). Thus, controlling and reducing pollution is an effective strategy for aging and disease prevention.

## Materials and methods

### Worm strains, cultivation, and exposure to pollutants

*C. elegans* were cultured at 20°C with *E. coli* OP50 as food source on standard NGM plates or in S-Medium as indicated. For all experiments, nematodes were synchronized by isolating eggs with hypochlorite/NaOH. For lifespans and aged populations, nematodes were cultured with supplemental FUdR from L4 larval stage to maintain synchronization. Nematodes were exposed on adult day 1 (24h after L4) to 0.02 or 1.25 mg/mL 50 nm silica NPs (Kisker, Steinfurt, Germany) or 10, 25, or 50 μM iHg, e.g., mercuric chloride (HgCl_2_). Silica NPs properties were previously analyzed by dynamic light scattering and live cell imaging and described (Hemmerich and von Mikecz, 2013; Scharf et al., 2013). The following strains were provided by the Caenorhabditis Genetics Center (funded by NIH Office of Research Infrastructure Programs (P40 OD010440)) and used where indicated: Wild type N2; DR1572 daf-2(1368); CF1038 daf-16(mu86); AM140 rmIs132 [unc-54 p::Q35::YFP] (Morley et al., 2002).

### Life span assays

Life span assays were performed as previously described (Petrascheck et al., 2007; Piechulek & von Mikecz, 2018). Briefly, approximately ten age-synchronized *C. elegans* were seeded as L1/L2 larvae in 96-well plates containing S-media and 6 mg/ mL *E. coli* OP50. To maintain worms age-synchronized, FUdR was added (1.5 mM final concentration, Sigma) at the L4 larval stage. On adult day 1, worms were left untreated or treated with particles or mercury at concentrations as indicated. Living animals were scored every second day with a dissecting scope. 67–262 worms were tested per group per experiment. Weibull plot was applied to the data of the survival analyses, e.g. curve fitting of the Kaplan-Meier survival curves.

### Mass spectrometry (LC-ESI-MS)

Characterization of *C. elegans* aggregomes were performed as previously reported (Arnhold et al., 2015; Scharf et al., 2016). Exposed nematodes were lysed in SDS-lysis buffer (150mM NaCl, 10mM Tris-HCL pH8, 2% SDS, protease inhibitor) and SDS-insoluble proteins were separated from the soluble protein fraction via filter retardation assay (Scharf et al., 2013). The SDS-insoluble protein fraction was eluted from cellulose acetate membranes with 6M guanidinium hydrochloride (GuaHCl). Dithiothreitol (DTT) was added to the GuaHCl solution to a final concentration of 10 mM and samples were incubated for one hour at room temperature. After addition of iodoacetamide to a final concentration of 20 mM and incubation for one hour at room temperature in the dark, samples were equilibrated to trypsin digestion condition by multiple cycles of ultrafiltration (MWCO=5000 Da, Microcon, Millipore) and buffer exchange against trypsin digestion buffer (8% acetonitrile in 25 mM ammonium bicarbonate). The sample solutions were digested with 15 ng trypsin overnight, dried in a rotary evaporator, dissolved in 20 μl 0.1% formic acid (FA) in 5% acetonitrile, centrifuged for 2 min (14,000 rpm) and the supernatants were transferred to a polypropylene-sample vial. Five μl of each sample were analyzed by MS and where necessary the remaining 15 μl were used for one additional measurement.

For mass spectrometry, the LTQ Orbitrap XL ETD (Thermo Scientific) coupled to a nano-HPLC NanoLC 2D System AS 1 (Eksigent) was used. The samples were loaded onto a trap column (Robust Reversed Phase Solid Phase Extraction Trap 100 μm × 40 mm, NanoSeparations) and washed with 30 μl buffer A (5% acetonitrile/0.1% formic acid). The bound analytes were transferred to a separation column (75 μm×100 mm,NanoSeparations) by applying a linear gradient from 0 to 38% of buffer B (80% acetonitrile/0.1% formic acid) over 76 min.

The measured spectra (DDA, top 8, ion trap) were processed by ProteomeDiscoverer 1.4 (ThermoScientific) and searched against the SwissProt (December 2015) database by Mascot (MatrixScience). The following search parameters were used in all searches: enzyme - trypsin with two allowed miss cleavages; fixed modification - carbamidomethylation of cysteine; variable modification - oxidation of methionine, phosphorylation of serine and threonine; measurement precision of precursor ions - 10 ppm; measurement precision of fragment ions - 0.5 Da.

Data compilation was performed by Scaffold 4 (Proteome Software) using the Mascot dat files and the msf files of ProteomeDiscoverer 1.4 of the measurements. The following parameters were used for the evaluation by Scaffold: Protein Grouping Strategy: Experiment-wide grouping with binary peptide-protein weights, Peptide Thresholds: 95,0% minimum, Protein Thresholds: 99,0% minimum and 2 peptides minimum, Peptide FDR: 0,0% (Decoy), Protein FDR: 0,0% (Decoy).

### Aggregome analysis

For the age-induced agregome, proteins that were identified in 2 out of 4 biological replicates were subjected to further analysis (Table S3), whereas for the iHg-induced aggregome, proteins identified in the 3 biological were pooled (Table S6). Identification and analysis of candidates for the silica NPs-induced aggregome were previously reported (Scharf et al., 2016). For further characterization, the identified aggregome candidates were analyzed by data mining of the following databases: PANTHER (Mi et al., 2013), UniProt (Bateman et al., 2021), and WormBase (Harris et al., 2020), and MalaCards (Rappaport et al., 2017). Specifically, each identified insoluble protein was categorized according to its biological process classified using the PANTHER database, UniProt, WormBase, and literature. Structural protein feature were analyzed as previously described (Arnhold et al., 2015). Briefly, structural feature information for each aggregome network were extracted from the UniProt database and percentage of aggregome network proteins with beta-strands, coiled-coil, cross-link, DNA binding, disulfide bonds, helices, metal binding feature, nucleotide binding feature, repeats, and zinc fingers was calculated, respectively. Then, statistical over- or underrepresentation was calculated by comparing to the average percentage of proteins in the total *C. elegans* genome with each feature.

For the comparative MS data analysis of aged aggregomes, SDS-insoluble data sets from two previously publications (David et al., 2010; Reis-Rodrigues et al., 2012) were compared to the identified age-induced aggregome network of this study (Table S4,5). In addition, the proteomic data set published in Walther et al., (2015) were used for analysis of protein abundances.

### Microscopy and Quantification of polyQ aggregates

*C. elegans* expressing repeats of 35 glutamines fused to yellow fluorescent protein in body wall muscle cells (AM140 rmIs132 [unc-54 p::Q35::YFP] (Morley et al., 2002)) were exposed to iHg or mock(H2O)-treated on adult day 1. After 24 hours, living hermaphrodites were imaged on 5% agarose pads/10 μM NaN3 with a 60x/1.4NA Plan Apo objective. Fluorescence quantifications was performed as described previously (Scharf et al., 2016).

### Statistical Analysis

Student’s t test was used to determine statistical significance with a p value below 0.05. The two parameters for the Weibull distribution were estimated to fit median survival function.

## Acknowledgments

We thank the Caenorhabditis Genetics Center (**funded by NIH Office of Research Infrastructure Programs (P40 OD010440))** for providing strains; WormBase; Kerry Kornfeld, Franziska Pohl, Kevin Nogutchi, and the Kornfeld lab for scientific input and discussion; Luke Schneider for experimental support; Jakob Risch for mathematical support. This work was funded by the Deutsche Forschungsgemeinschaft, Grant MI 486/10-1.

## Author Contributions

A.S. and A.vM. conceived and designed the experiments. A.S., A.P., and KH.G. performed experiments, A.S., A.P., and KH.G. analyzed data, A.S, A.P., KH.G. and A.vM provided scientific input, A.S, and A.vM wrote the paper.

