## Supplementary Figures for "Pollutants corrupt resilience pathways of aging in the nematode *C. elegans*"

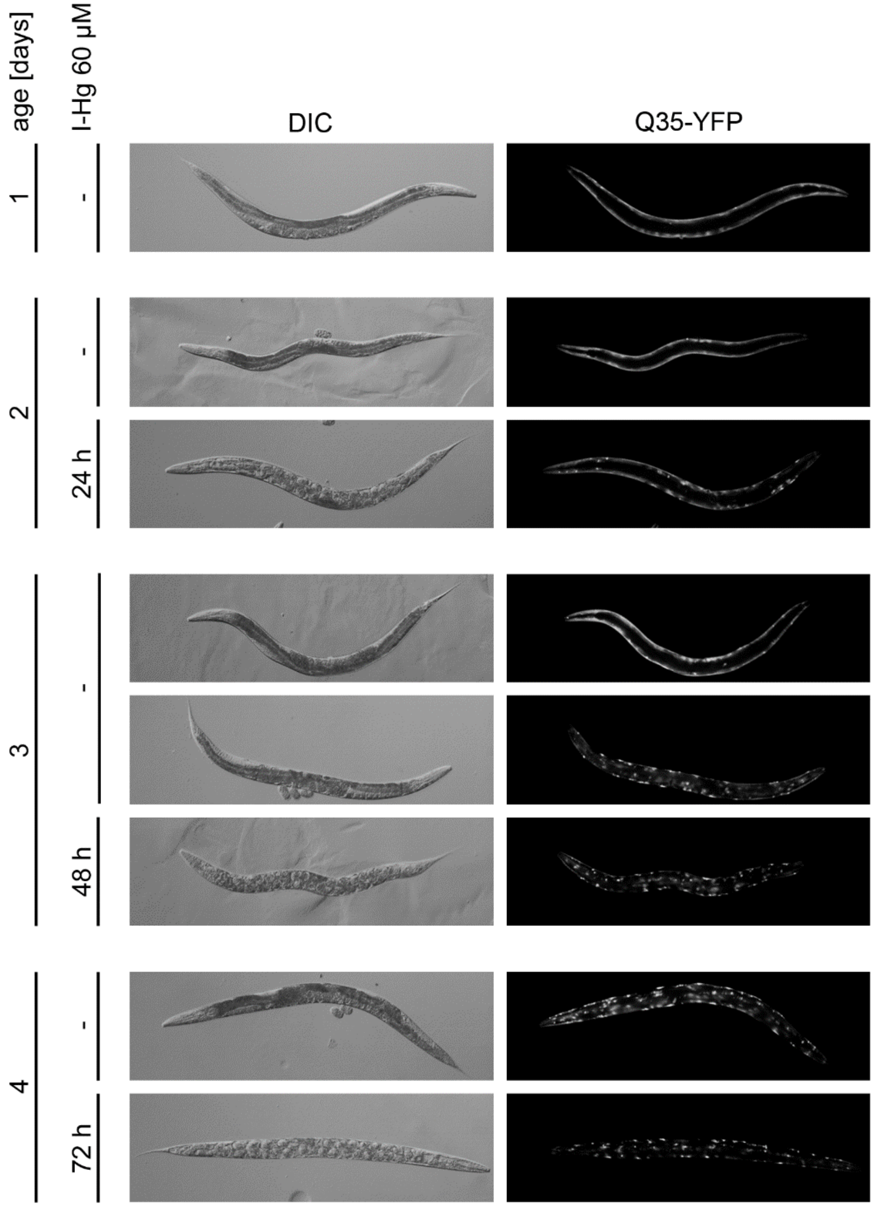


**Figure S1. iHg accelerates proteotoxic stress**. Representative fluorescent micrographs of 1, 2, 3, and 4 day old, adult hermaphrodites of a *C. elegans* proteotoxic stress model that expresses homopolymeric repeats of 35 glutamines fused to yellow fluorescent protein (Q35::YFP) in body wall muscle cells (Morley et al., 2002). Hermaphrodites were mock-treated or exposed to 50 µM iHg for 24, 48, and 72 hours. Bar, 100 µM. Mock-treated showed an age-related change from a mostly smooth distribution of yellow fluorescence along the body wall muscles to a dotted pattern of aggregated polyQ. Hermaphrodites exposed to iHg exhibit an acceleration of the age-related change to aggregated polyQ with a first appearance at adult day 2. Note that the two depicted untreated hermaphrodites for the 48-hour time-point reflect the wide range of aggregation progression observed in this cohort.

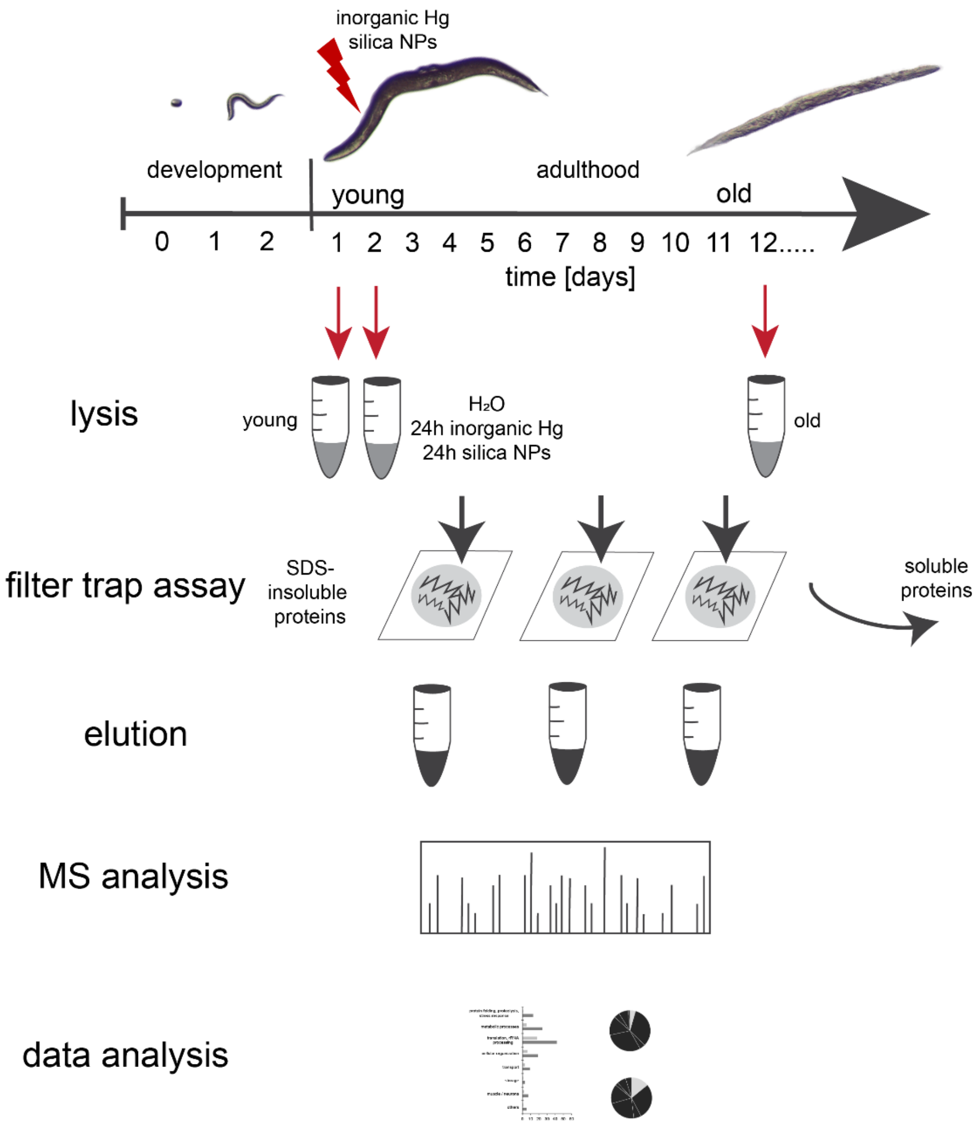


**Figure S2. Schematic of the experimental design of the different aggregome characterizations**. For the iHg- and silica NPs- induced aggregome analysis, synchronized wild type *C. elegans* (N2) were mock-treated, or exposed to iHg or silica NPs on adult day 1 for 24 hours before lysis. For the age-induced aggregome analysis, synchronized wild type *C. elegans* (N2) were lysed on adult day 1 as “young” or on adult day 12 as “old”. SDS-insoluble proteins were trapped on filter, eluted with 6M guanidinium hydrochloride, followed by mass spectrometry (ESI-LC/MS), and data analysis including data mining. Note that the experiments to identify the iHg, the silica NPs, and the age-induced aggregome networks were done separately.

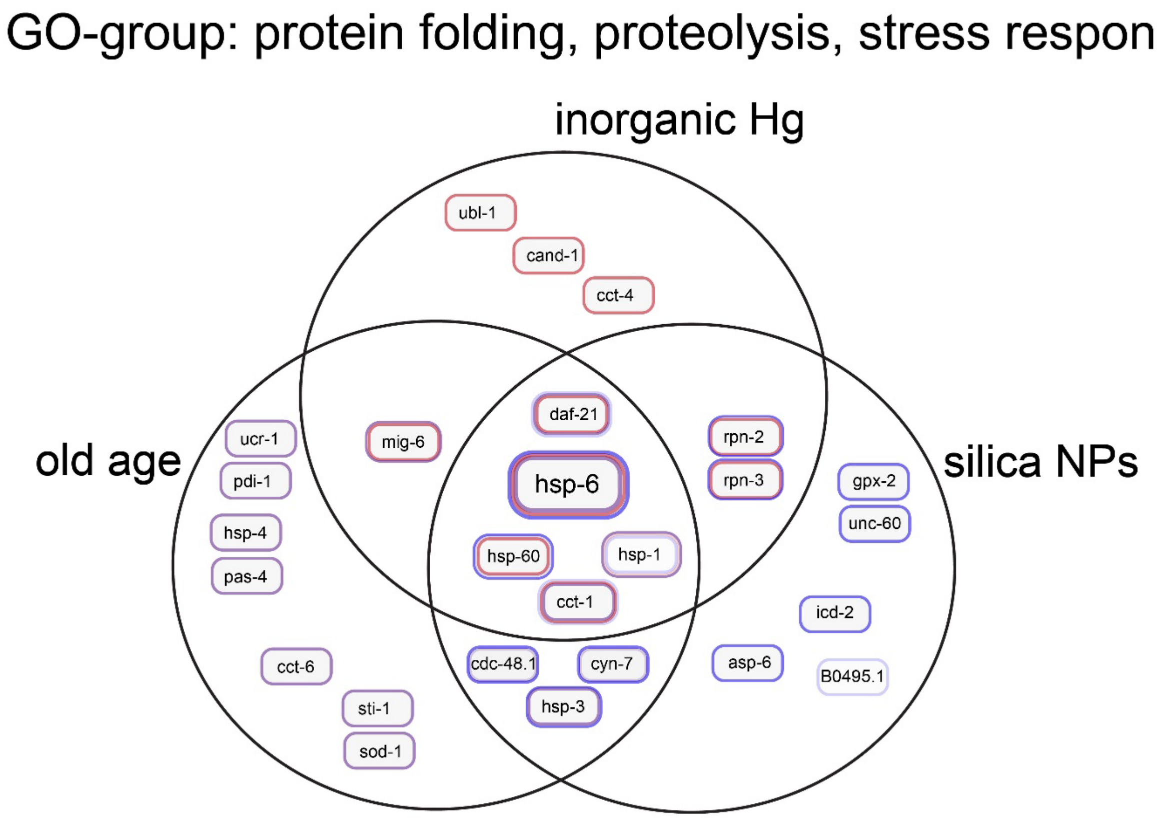


**Figure S3. GO-group: protein folding, proteolysis, and stress response**. Venn diagram of identified insoluble proteins categorized according their biological process protein folding, proteolysis, and stress response classified by the PANTHER databank (Mi et al., 2013) in the iHg- (red), silica NPs- (blue, (Scharf et al., 2016)), and age-induced (purple) aggregome network. The boxes indicate the number of proteins identified in the ground and specific aggregome, respectively.

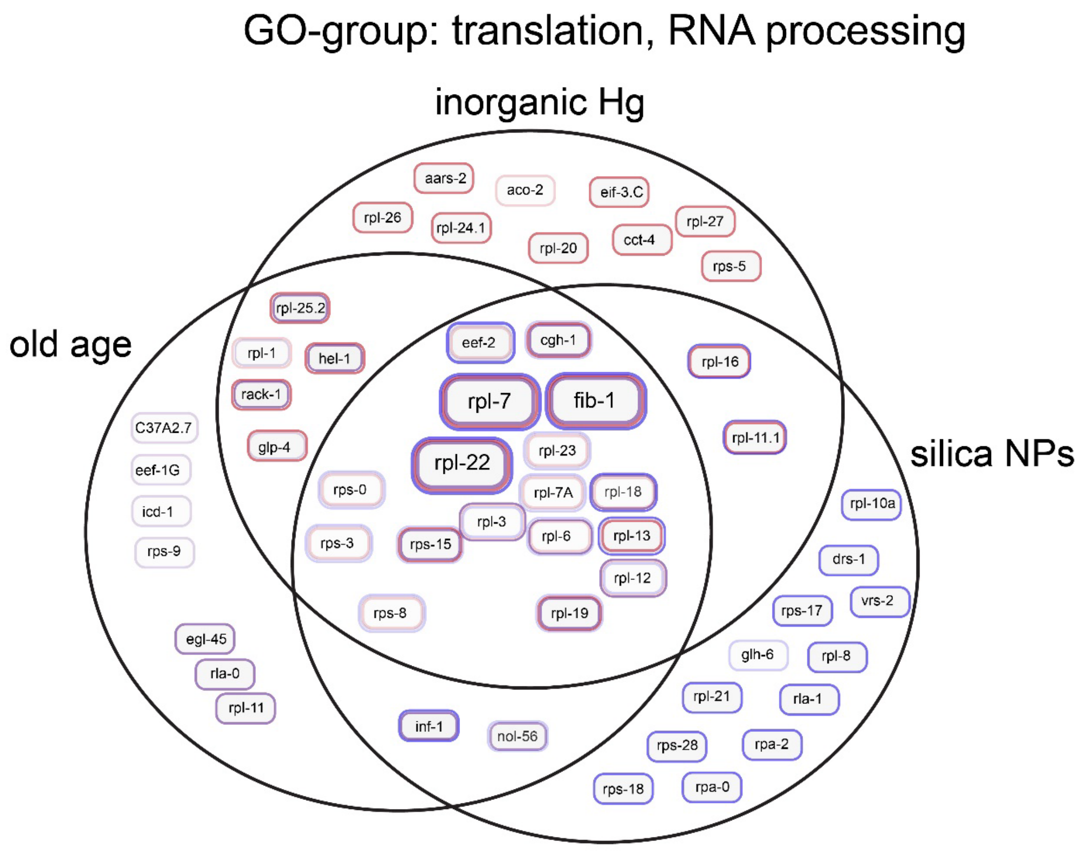


**Figure S4. GO-group: translation and RNA pocessing**. Venn diagram of identified insoluble proteins categorized according their biological process translation and RNA processing classified by the PANTHER databank (Mi et al., 2013) in the iHg-(red), silica NPs- (blue), and age-induced (purple) aggregome network.

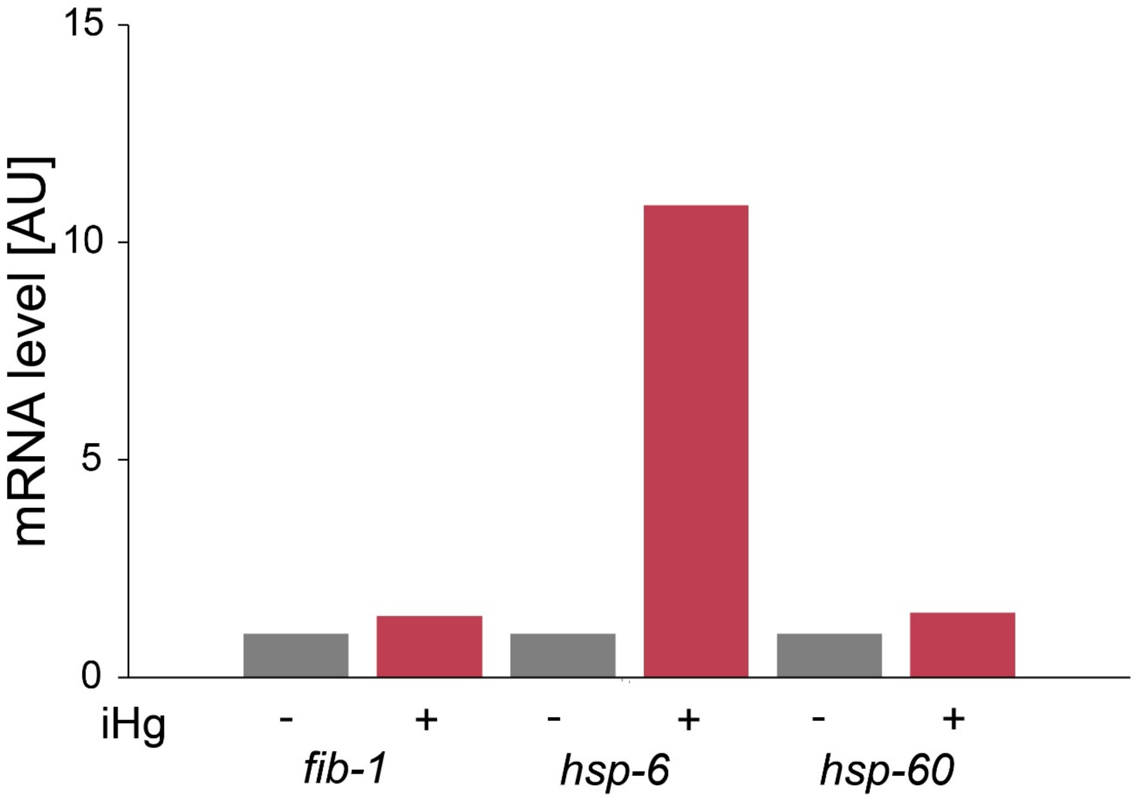


**Figure S5. Expression is not correlated with protein aggregation**. The bar graph shows mRNA levels of *fib-1*, *hsp-6*, and *hsp-60* in $H_2O$-treated versus 50 μM iHg-treated wild type C. *elegans*. RNA levels were determined by qPCR in two independent experiments and show fold change compared to $H_2O$ control.
