## Supplementary Tables S1 S2 for "Pollutants corrupt resilience pathways of aging in the nematode *C. elegans*"

**Supplementary Table 1**. Lifespan summary statistics. Related to Figure 1.

| Strain | Treatment[1] | Mean adult lifespan +/- SD (days)[2] | Maximum adult lifespan (days)[3] | N[4](n[5]) |
|---|---|---|---|---|
| N2 | $HgCl_2$ 0 µM | 22 | 32 | 2(144) |
| N2 | $HgCl_2$ 10 µM | 19 | 27 | 2(167) |
| N2 | $HgCl_2$ 25 µM | 8 | 12 | 2(171) |
| N2 | $HgCl_2$ 50 µM | 3 | 7 | 2(168) |
| N2 | silica NPs 0 mg/ml | 23 +/-2 | 37 +/-3 | 3(406) |
| N2 | silica NPs 0.02 mg/ml | 23 +/-2 | 37 +/-2 | 3(395) |
| N2 | silica NPs 1.25 mg/ml | 15 +/-3 | 28 +/-3 | 3(454) |
| Daf-2 | silica NPs 0 mg/ml | 31 +/-2 | 51 +/-4 | 3(344) |
| Daf-2 | silica NPs 0.02 mg/ml | 31 +/-3 | 50 +/-4 | 3(344) |
| Daf-2 | silica NPs 1.25 mg/ml | 26 +/-2 | 40 +/-3 | 3(320) |
| Daf-16 | silica NPs 0 mg/ml | 16 +/-4 | 31 +/-1 | 3(426) |
| Daf-16 | silica NPs 0.02 mg/ml | 12 +/-4 | 29 +/-2 | 3(414) |
| Daf-16 | silica NPs 1.25 mg/ml | 16 +/-5 | 21 +/-4 | 3(416) |

Values are mean with standard deviation (SD).
[1]treatment:
[2]mean adult lifespan: average adult lifespan.
[3]maximum adult lifespan: lifespan of the longest lived worm.
[4]N: Number of independent laboratory experiments or simulations.
[5]n: Number of worms analyzed.

**Supplementary Table S2.** Silica NPs and iHg induce protein fibrillation.

| Experimental system | Phenotype | Silica NPs | iHg |
|---|---|---|---|
| **human cells[1]** | global amyloid fibrillation | ND | yes (Arnhold et al., 2015) |
| | local amyloid fibrillation | yes, nucleus (Chen et al., 2008) | yes: nucleus, nucleolus (Arnhold et al., 2015) |
| ***C. elegans*[2]** | global amyloid fibrillation | yes (Scharf et al., 2013, 2016) | yes, this study |
| | local amyloid fibrillation | yes, intestinal nucleolus (Scharf et al., 2016) | yes, intestinal nucleolus (Arnhold et al., 2015) |

[1]human cells: human cell culture with HEp-2 or SH-SY5Y

[2]*C. elegans* wild type N2

ND, not determined
